# Cell type-specific aging clocks to quantify aging and rejuvenation in regenerative regions of the brain

**DOI:** 10.1101/2022.01.10.475747

**Authors:** Matthew T. Buckley, Eric Sun, Benson M. George, Ling Liu, Nicholas Schaum, Lucy Xu, Jaime M. Reyes, Margaret A. Goodell, Irving L. Weissman, Tony Wyss-Coray, Thomas A. Rando, Anne Brunet

**Author notes:** These authors contributed equally (Matthew T. Buckley, Eric Sun).

## Abstract

Aging manifests as progressive dysfunction culminating in death. The diversity of cell types is a challenge to the precise quantification of aging and its reversal. Here we develop a suite of ‘aging clocks’ based on single cell transcriptomic data to characterize cell type-specific aging and rejuvenation strategies. The subventricular zone (SVZ) neurogenic region contains many cell types and provides an excellent system to study cell-level tissue aging and regeneration. We generated 21,458 single-cell transcriptomes from the neurogenic regions of 28 mice, tiling ages from young to old. With these data, we trained a suite of single cell-based regression models (aging clocks) to predict both chronological age (passage of time) and biological age (fitness, in this case the proliferative capacity of the neurogenic region). Both types of clocks perform well on independent cohorts of mice. Genes underlying the single cell-based aging clocks are mostly cell-type specific, but also include a few shared genes in the interferon and lipid metabolism pathways. We used these single cell-based aging clocks to measure transcriptomic rejuvenation, by generating single cell RNA-seq datasets of SVZ neurogenic regions for two interventions – heterochronic parabiosis (young blood) and exercise. Interestingly, the use of aging clocks reveals that both heterochronic parabiosis and exercise reverse transcriptomic aging in the niche, but in different ways across cell types and genes. This study represents the first development of high-resolution aging clocks from single cell transcriptomic data and demonstrates their application to quantify transcriptomic rejuvenation.

## Introduction

Aging is the progressive deterioration of cellular and organismal function. Age-dependent decline is linked in large part to the passage of time and therefore the chronological age of an individual. But such chronological decline is not inexorable. At the same chronological age, some individuals have better organismal and tissue fitness (biological age) than others. Furthermore, aging trajectories can be slowed, and some aspects of aging can be reversed by specific interventions, including dietary restriction, exercise, senolytics (compounds that kill senescent cells), reprogramming factors, and young blood factors^1–6^. As aging is the primary risk factor for many diseases, particularly neurodegenerative diseases^7–10^, a better understanding of aging and ‘rejuvenation’ strategies could yield large benefits for a wide-range of diseases.

Aging is complex and difficult to quantify. One quantification approach is to use machine learning to build age-prediction models – aging clocks – which can serve as integrative aging biomarkers. Such clocks should also accelerate our understanding of existing interventions and help identify new strategies to counter aging and age-related diseases. Machine learning models trained on high dimensional datasets (e.g. DNA methylation, transcriptomics, proteomics) have been shown to predict chronological age with remarkable accuracy. For example, regression-based aging clocks trained on DNA methylation profiles from multiple tissues (‘epigenetic aging clocks’)^11–15^ or blood plasma protein profiles^16–19^ have striking performance to predict chronological age in humans. Aging clocks directly optimized to predict biological age have also been developed by regressing functional phenotypes^14,15^ or time remaining until death^20,21^. Beneficial health interventions such as diet and exercise^22–24^ and genetic manipulations^25–27^ result in younger predictions from epigenetic aging clocks trained on chronological age. Hence, epigenetic aging clocks, despite being trained on chronological age, also capture dimensions of biological age.

So far, molecular aging clocks have largely relied on datasets built using bulk tissue input or purified cell populations^11–15,28–33^. Bulk tissue profiles (and even purified populations) average the molecular profiles from many cells, integrating tissue composition changes and cell type-specific responses. Hence, the cell type-specific contributions to aging and rejuvenation detected by these clocks remain unclear. While single cell DNA methylation and transcriptomic have started to be used to classify age^34,35^, cell-specific transcriptomic aging clocks have not yet been generated. Thus, it remains to be determined if different cell types’ clocks ‘tick’ at different rates, which cell types predict age most accurately, and how specific cell types respond to different interventions. The rapid advance of single cell RNA-sequencing technologies provides a unique opportunity to explore these unaddressed questions and identify new molecular aging clocks to study interventions to counter aging and age-related diseases.

## Results

### Cell type-specific transcriptomic aging clocks

As a paradigm for tissue aging and functional decline in the brain, we focused on the neurogenic region located in the subventricular zone (SVZ) of the adult mammalian brain. The SVZ neurogenic region contains neural stem cells (NSCs) that give rise to differentiated cells (neurons, astrocytes) that are important for olfactory discrimination and repair upon injury^36–43^. Importantly, this neurogenic region contains at least 11 different cell types and experiences age-related changes correlated with deterioration in tissue function^40,44–47^. We built cell type-specific aging clocks trained to predict the chronological or biological age of the subventricular zone neurogenic niche. To train these clocks, we performed single cell RNA-seq on neurogenic regions from 28 mice, tiling 26 different ages from 3 months (young adult) to 29 months (geriatric adult) (Fig. 1a). Given the constraining cost of single-cell RNA-seq, we used lipid-modified oligonucleotide labeling (MULTI-seq, see Methods)^48^ to multiplex samples within four independent cohorts, each with 4-8 mice (Supplementary Table 1). After demultiplexing and quality control, we obtained 21,458 high quality single cell transcriptomes (Extended Data Fig. 1a, b). Clustering and UMAP visualization confirmed the presence of 11 cell types, including differentiated cell types (oligodendrocytes, microglia, endothelial cells) and cells from the neural stem cell lineage (astrocytes and quiescent NSCs (qNSCs), activated NSCs (aNSCs) and neural progenitor cells (NPCs), and neuroblasts) (Fig. 1b). This analysis also corroborated the decline of proliferating NSCs in this region during aging (Fig. 1c, d)^40^. Our dataset provides a high temporal resolution resource to characterize aging in a regenerative stem cell region in the brain.

**Figure 1:**
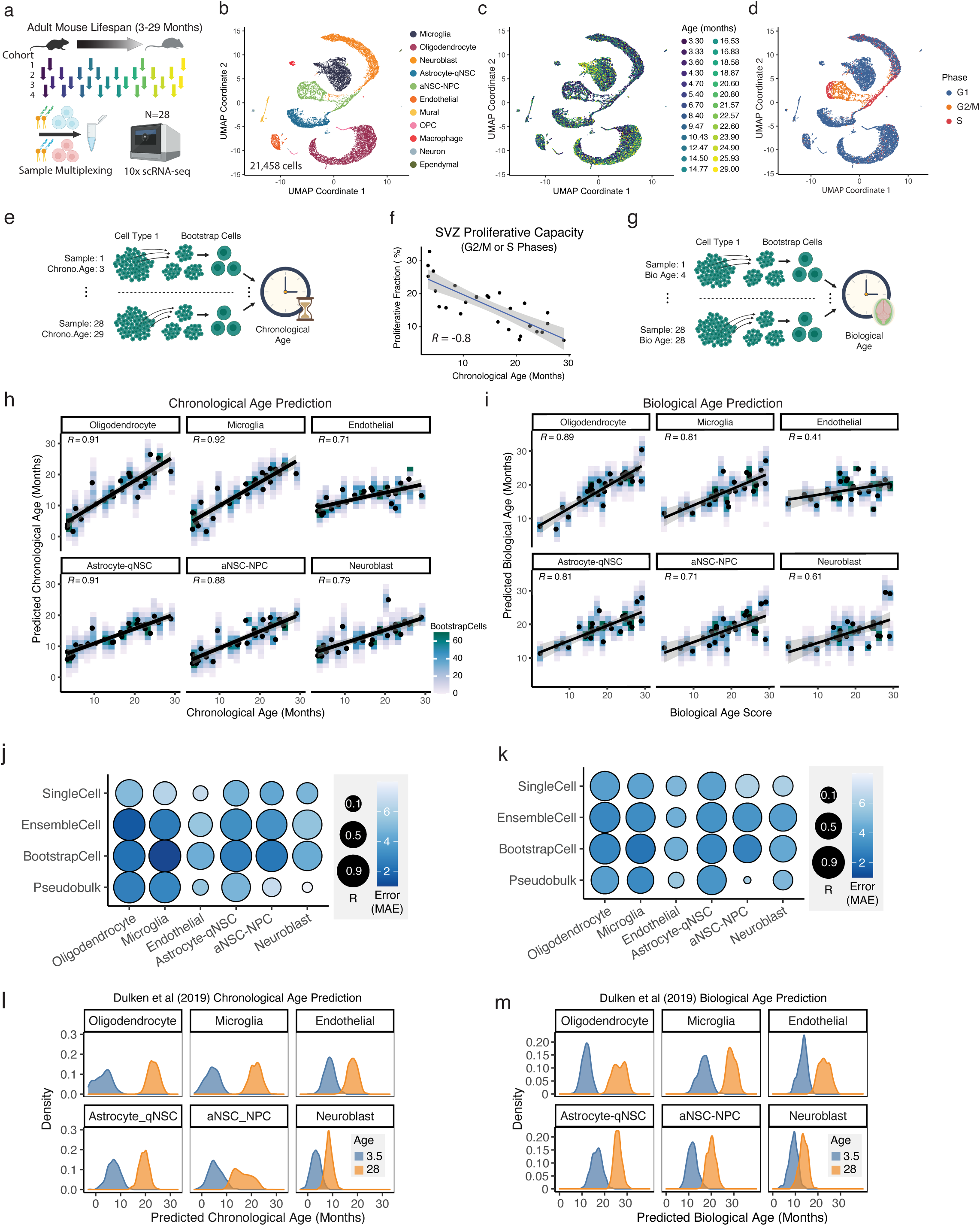
Cell-type specific transcriptomic aging clocks for neurogenic regions. **a,** Training data for single-cell transcriptomic aging clocks. 10x Genomics single cell transcriptomics was performed on microdissected subventricular zone (SVZ) neurogenic regions from four independent cohorts of 4-8 from male mice, aged 3.3 to 29 months (see Supplementary Table 1). SVZs from the same cohort were multiplexed using lipid-modified oligonucleotide labeling (MULTI-seq). **b,** UMAP projection and cell type clustering of SVZ single cell transcriptomes integrated across cohorts. 21,458 cells passed quality control and doublet filtering. 11 cell types were detected. Each dot represents the transcriptome of an individual cell with transcripts detected from at least 500 genes. **c,** The same projection of 21,458 transcriptomes from (b) but colored by the age of the mouse from which the cell was sampled. Note that two pairs of mice had the same age, resulting in 26 different age colors corresponding to 28 mice. **d,** The same projection of 21,458 transcriptomes from (b) but colored by the predicted cell cycle state based on Seurat’s CellCycleScoring function with the default cell cycle markers. Activated neural stem cells and neural progenitor cells (aNSC-NPCs) represent the vast majority of cells predicted to be in G2/M and S phases in the SVZ neurogenic niche. **e,** Schematic depicting the process of generating BootstrapCells for training chronological clocks. From a population of cells from given cell type and sample, 15 cells are sampled without replacement and combined to generate one BootstrapCell. This process is repeated 100 times per cell type-sample combination, to generate a training dataset that equally weighs each SVZ sample. **f,** SVZ proliferative fraction (cells predicted to be G2/M or S phase) as function of chronological age, quantifying the relationship between (c) and (d). R represents Pearson’s correlation coefficient. **g,** Schematic depicting the process of generating BootstrapCells for training biological age clocks. Biological age is defined as a function of the proliferative capacity of the SVZ rather than chronological age. **h,** Performance of BootstrapCell chronological age prediction across cell types. Density of BootstrapCell predictions is depicted in color and overlaid black dots represent the median prediction for each sample. Performance is based on cross-cohort-validation. R values are Pearson’s correlation coefficients at the sample level. **i,** As in (h) but for BootstrapCell biological age score prediction across cell types. Biological age score is a linear transformation of the SVZ proliferative fraction to rescale the range to be comparable (but not identical) to chronological age in months. **j,** Overview of Pearson’s correlation coefficients and median absolute error values for various methods of predicting chronological age across cell types from single-cell transcriptomic data. SingleCell uses *bona fide* single cell transcriptomes with minimal processing as input to a lasso regression model to predict chronological age. BootstrapCell uses a lasso model and the preprocessing method depicted in (e). EnsembleCell involves repeatedly partitioning cells into groups of 15 cells and training an ensemble of elastic net models to predict age. Pseudobulk involves naïve pseudobulking all cells from the same cell type and sample before using a lasso regression model to predict age. Performance is based on cross-cohort-validation. **k,** As in (j) but evaluating biological age prediction models rather than chronological age prediction models. **l,** External validation of BootstrapCell chronological age prediction models (chronological aging clocks) on single cell transcriptomic data from young and old SVZ samples by Dulken et al, 2019. Histograms show separated age prediction distributions, indicating ability to discriminate age. **m,** As in (l) but evaluating biological age prediction models (biological aging clocks).

To develop robust single cell-based aging clocks, we focused on the six most abundant recovered cell types in the SVZ – oligodendrocytes, microglia, endothelial cells, astrocytes-qNSCs (which cluster together, see Fig. 1b), aNSC-NPCs (which cluster together, see Fig. 1b), and neuroblasts. We first developed chronological age models that maximize correlation and minimize error between predicted and true chronological age. We built different models (lasso and elastic net regression^49–51^) from single cell transcriptomic data for each of the 6 cell types as an input (see Methods). We evaluated performance of the models on true chronological age, by building models on 3 of the 4 cohorts and testing it on the remaining cohort (cross-cohort-validation). This strategy avoids performance inflation caused by training and evaluating on correlated cells from the same animal or animals from the same cohort. Our resulting top performing chronological aging clocks, termed ‘Bootstrap’ and ‘Ensemble’, are groups of lasso and elastic net models trained on either bootstrap-sampled or randomly partitioned and merged meta-cells, termed BootstrapCells or EnsembleCells (see Methods). For the Bootstrap models, 100 BootstrapCells are generated by taking 100 random samples of 15 cells for each cell type and animal combination, such that each animal contributes equally to the training data (Fig. 1e). For the Ensemble model, a random partitioning and elastic net model training process is repeated 100 times then combined to generate a single Ensemble model for a given cell type, such that each cell contributes equally. In our cross-cohort validation, these two models performed well to predict chronological age. For example, oligodendrocyte Bootstrap models predicted true chronological age most successfully with a correlation (R) = 0.91 and an error = 1.6 months and microglia Bootstrap models predicted chronological age with R = 0.92 and an error = 2.1 months (Fig. 1h, Supplementary Table 2). Such performance in a cross-cohort-validation scheme suggests that these chronological aging clocks are not batch dependent. Overall, these models had R ranging from 0.71-0.92 and errors ranging from 1.6-5.4 months, and they uniformly surpassed the performance of raw single-cell trained clocks and pseudo-bulked clocks (i.e. pool of all cells from each cell type) (Fig. 1j, Extended Data Fig. 1c, Supplementary Table 2).

To externally validate these chronological aging clocks, we retrained clocks on all 28 mice and applied them to independent datasets. Our single cell-based models easily separated young and old samples in an independent single cell RNA-seq dataset from SVZ neurogenic regions of young and old mice^46^ (Fig. 1l). The predicted age in months was more accurate with some cell types (e.g. microglia) than others (e.g. neuroblasts) (Fig. 1l). These aging clocks also correctly ordered RNA-seq samples from another neurogenic region in the brain – dentate gyrus of hippocampus^52^, properly discriminating mice only a month apart in age (Extended Data Fig. 1d). The successful application of our chronological aging clocks to these independent datasets demonstrates their robustness and generalizability to other regenerative brain regions.

We also developed biological aging clocks from our single cell transcriptomic data. While chronological aging clocks can record aspects of biological age^17,23,25^ and predict disease probability^53,54^, clocks trained on functional metrics linked to biological age may be particularly useful for intervention assessment^14,25^. The primary functional role of the SVZ neurogenic region is to harbor proliferating NSCs that can produce new neurons which in turn integrate into functional neural circuits^40–43^. The proliferative capacity of NSCs in the SVZ neurogenic region declines with age, and this decline may be considered a functional metric of biological age of this region^55–58^. To define neurogenic region ‘fitness’, we quantified the proliferative fraction of cells (consisting almost exclusively of aNSC-NPCs and neuroblasts) in the neurogenic regions from each of the 28 mice, based on cell cycle signatures (see Fig. 1d). The fraction of cells predicted to be proliferative (in G2/M and S phases; the ‘proliferative fraction’) ranged from 5-30% and was negatively correlated with chronological age, as expected (*R = -*0.8) (Fig. 1f). Although measures of proliferative fraction are subject to technical noise, they may still be helpful to capture biological age. We therefore trained a suite of clocks analogous to those described above, except using aNSC-NPC ‘proliferative fraction’ as a proxy for biological age instead of chronological age (Fig. 1g). These biological aging clocks achieved robust prediction performance, though slightly diminished in comparison to chronological aging clocks (R = 0.41-0.89, error = 2.3 - 4.6 months) (Fig. 1i, k). They also separated samples by age in external datasets (Fig. 1m, Extended Data Fig. 1e). Even though all clocks were trained on aNSC-NPC ‘proliferative fraction’, the microglia and oligodendrocytes biological age clocks performed better than aNSC-NPC ones. Collectively, these data reveal that single cell transcriptomes can be used to build accurate chronological and biological aging clocks for different cell types.

### Genes that contribute to the cell-type specific aging clocks

What makes these aging clocks ‘tick’ – i.e. what are the genes that contribute to the aging clocks in each cell type? In the process of training, each clock selects top genes useful for accurate prediction and weighs the importance of each. To analyze selected genes and their relative contributions, we visualized each chronological aging clock as a donut plot with genes that contribute the most at the top (Fig. 2a, Extended Data Fig. 2a). Gene sets contributing to the chronological aging clocks in different cell types ranged from 96 to 359 genes, and they encompassed genes whose expression increased with age (orange) or decreased with age (blue) (Fig. 2a, Extended Data Fig. 2a, Supplementary Table 3). The top genes contributing to the aNSC-NPC chronological aging clock were *AC149090.1* and *Ifi27*, which are both upregulated with age (orange) (Fig. 2a). *AC149090.1* is orthologous to human *PISD*, a gene encoding a phospholipid decarboxylase involved in lipid metabolism (phosphatidylethanolamine production), linked to autophagy, and localized to the inner mitochondrial membrane^59,60^. *Ifi27* (also referred to as *Isg12)* is a transcript upregulated in response to type I interferons^61^ (Fig. 2a). Thus, aging clocks identify many genes, including inflammation and lipid metabolism genes, whose expression is most predictive of aging in a particular cell type.

**Figure 2:**
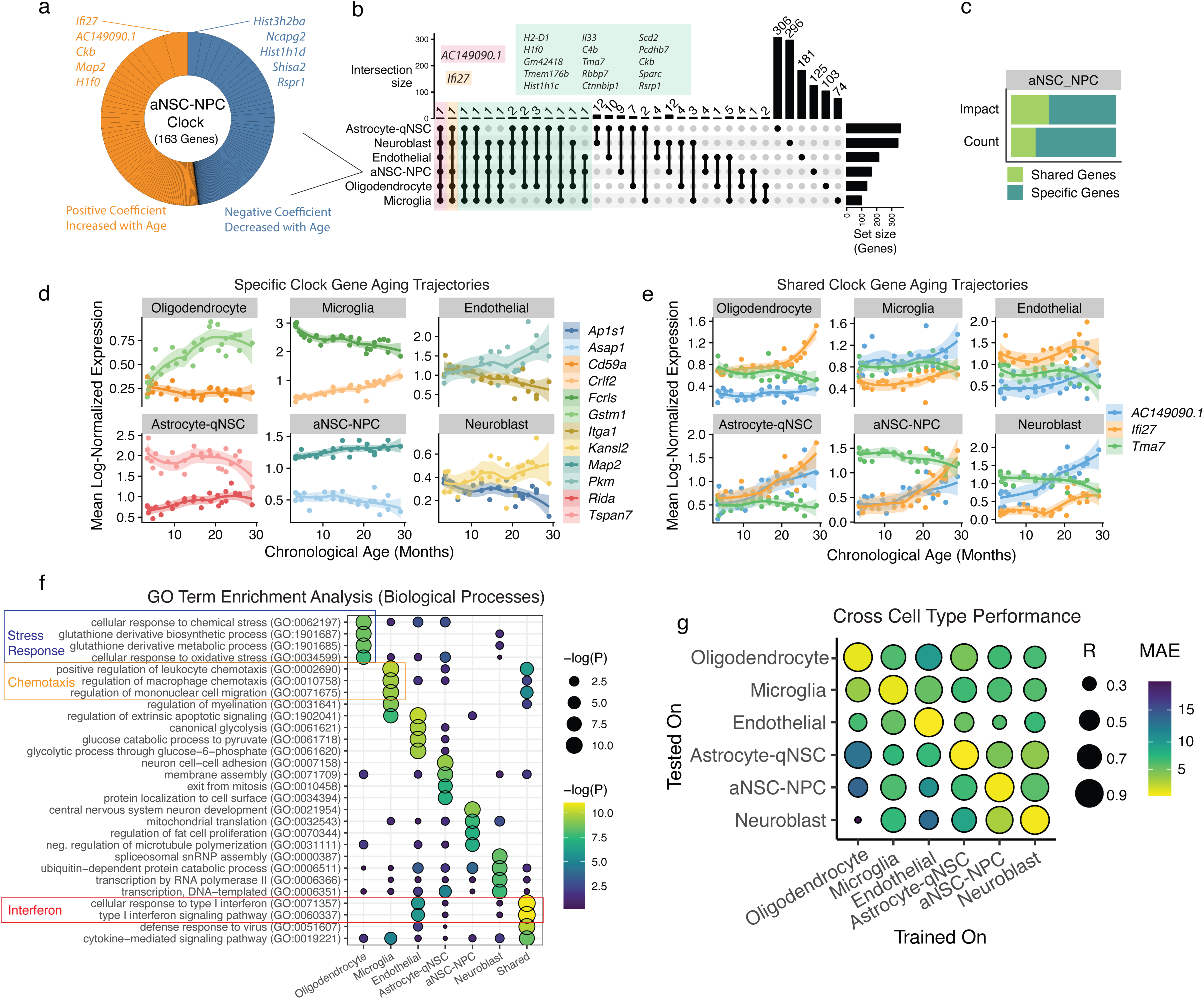
Genes underlying the cell-type specific chronological aging clocks. **a,** Contribution of individual genes to the aNSC-NPC chronological aging clock (BootstrapCell). Donut plots, with sector size denoting gene weight in the model and color indicating sign of expression change with age. Total number of genes used by the clock is provided in the center of each donut plot. Positive coefficients (orange) indicate higher gene expression is associated with older age. Negative coefficients (blue) indicate lower gene expression is associated with older age. For example, *Ifi27* has the largest positive coefficient – i.e. higher expression of *Ifi27* contributes to an older predicted age. For genes contributing to the other cell-specific chronological aging clocks, see Extended Data Fig. 2a. All genes and coefficients can be found in Supplementary Table 3. **b,** Upset plot illustrating the intersection of gene sets used by cell type-specific chronological aging clocks. Genes present in four, five, or six of the analyzed clocks are highlighted in green, yellow, and red, respectively. Only one gene, *AC149090.1*, is selected by all six clocks; the vast majority of clock genes are cell type-specific. For genes selected by biological aging clocks, see Extended Data Fig. 2b. **c,** Barplot comparing impact and count of shared and specific genes within the aNSC-NPC chronological aging clock. Impact is the sum of the absolute value of the gene coefficients. Count is the number of genes in each category. Shared genes are the minority but have an outsized influence. For other cell-specific chronological aging clocks, see Extended Data Fig. 2c. **d,** Expression trajectories as a function of age of select clock-specific genes. Expression values are log-normalized counts per 10,000 transcripts. **e,** Expression trajectories as a function of age of select shared genes across at least four cell type-specific clocks. **f,** Top enriched Gene Ontology (GO) Biological Process terms from gene set enrichment analysis of genes selected by chronological aging clocks. Shared genes (present in two or more clocks) are enriched for *cellular response to type I interferon* while cell type-specific gene sets show enrichment for largely unique terms, including *negative regulation of transcription* across neuroblast-specific clock genes and *positive regulation of cytokine secretion* in microglia-specific clock genes. **g,** Assessment of the ability of cell type-specific clocks to predict age given transcriptomes of different cell types. Size of dots corresponds to Pearson correlation with chronological age and color indicates median absolute error. Error substantially increases when testing on alternate cell types. For GO term analysis of cell-specific biological aging clocks, see Extended Data Fig. 2d.

To investigate whether each cell type-specific clock select similar or unique genes, we compared intersections of chronological aging clock gene sets (Fig. 2b, for genes contributing to biological aging clocks, see Extended Data Fig. 2b). Interestingly, *AC149090.1*, was selected by chronological aging clocks from all 6 different cell types (Fig. 2b) and another gene, *Ifi27*, was selected by chronological aging clocks from 5 out of 6 cell types (Fig. 2b). In contrast, most genes selected by the cell-type-specific clocks were cell type-specific (Fig. 2b). The specificity of the clocks exceeded what would be expected from transcriptome cell type specificity alone (Extended Data Fig. 3a, b). However, shared genes carry a disproportionate weight within the clocks, with coefficients approximately 40% larger in magnitude (Fig. 2c, Extended Data Fig. 2c). Cell type-specific genes (Fig. 2d) and even shared genes (Fig. 2e) exhibited differences in trajectory shapes (*Fcrls* and *Crlf2* in microglia) and expression magnitudes (e.g. *Ifi27* in different cell types from the NSC lineage) during aging in different cell types. Thus, cell type-specific clocks capture useful cell-type specific expression differences and dynamics that would be missed by bulk methods.

While genes selected by the aging clocks are mostly cell-type specific, the pathways to which they belong could still be widely shared across cell types. To test this possibility, we examined the pathways enriched in the specific or shared genes selected by the chronological aging clocks. Interestingly, gene set enrichment analysis (GSEA) on the specific genes from each clock revealed enrichment for different biological processes in each cell type (Fig. 2f, Extended Data Fig. 2d for biological aging clocks), e.g. stress response for oligodendrocytes and chemotaxis for microglia. Thus, pathways for specific genes selected by the clocks are also largely cell-type specific and may reflect age-dependent changes in function in each cell type. Even though there are some common genes across all clocks and these are more heavily weighted, cell type-specific clocks did not perform as well on other cell types (Fig. 2g). There were a few biological processes most enriched in shared genes, including response to type I interferon and cytokine signaling (Fig. 2f). Hence, our dissection of the genes composing the clocks highlights specific and common features of cellular aging, including stress response, lipid metabolism, and inflammation.

### Heterochronic parabiosis rejuvenates cell-specific aging clocks, notably activated neural stem cells clocks

Do single cell-based aging clocks – whether trained on chronological or biological age – capture known ‘rejuvenating’ interventions? A robust rejuvenating intervention across tissues is heterochronic parabiosis – the sharing of blood circulation between young and old animals^62–71^. Parabiosis with a young animal can restore aspects of cell function (e.g. NSC proliferation, vascular remodeling) in the SVZ neurogenic region of an old animal, and part of the effects can be recapitulated by the injection of young blood or plasma^65,72,73^. To test how our single cell-based aging clocks recorded the impact of exposure to young and old blood on regenerative regions, we generated multiplexed single-cell RNA-seq data on SVZ neurogenic regions from parabiosed young and old mice and isochronic parabiosed controls. In total, we collected 25,595 single cell transcriptomes from the SVZ neurogenic regions of 18 mice across 2 independent cohorts (Fig. 3a, Extended Data Fig. 4a, Supplementary Table 1) (See Methods). This dataset represents the first high quality single cell RNA-seq resource for heterochronic parabiosis in the SVZ neurogenic region.

**Figure 3:**
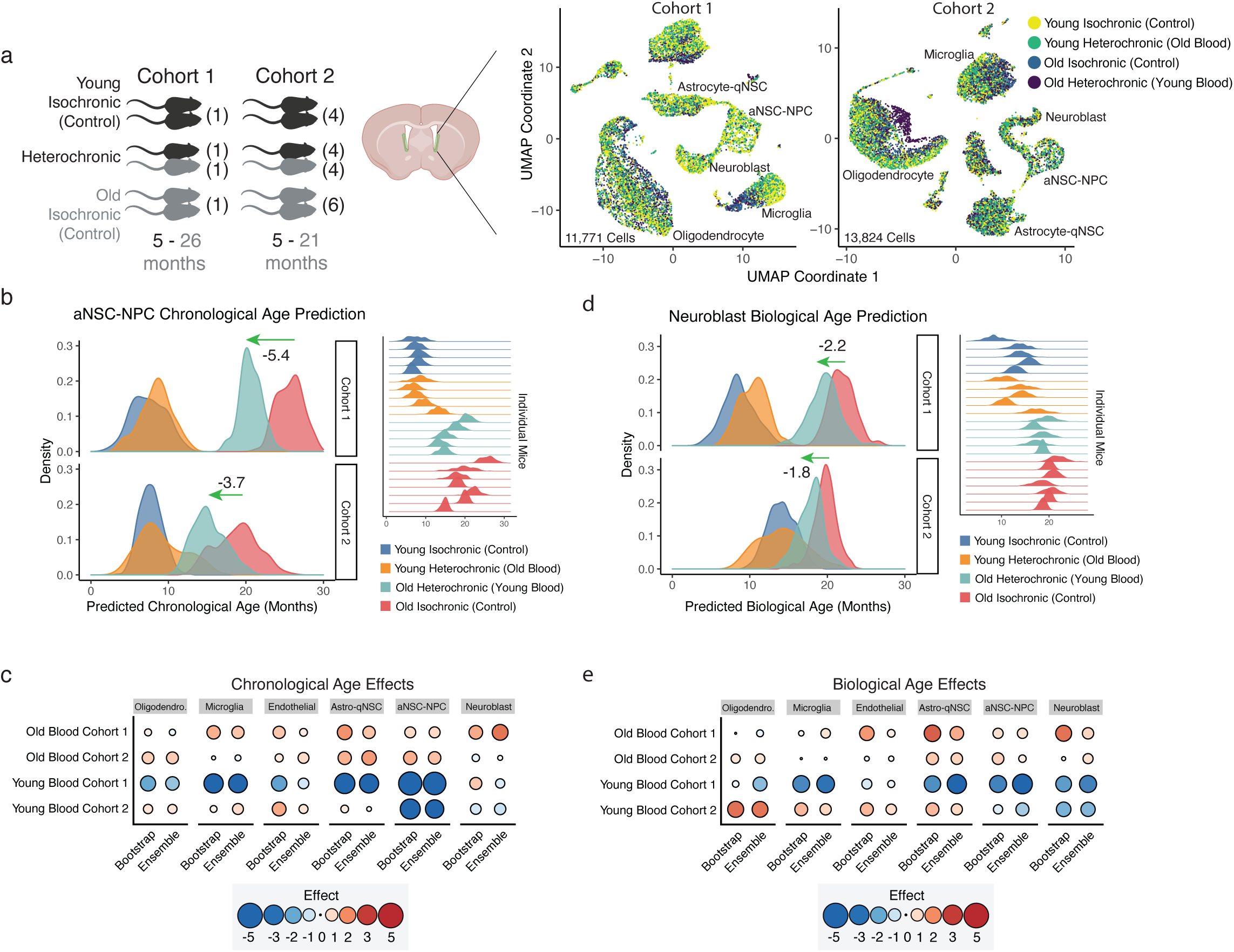
Effect of heterochronic parabiosis (young blood) on cell-type specific aging clocks. **a,** Schematic of parabiosis cohorts and corresponding UMAP projections from each cohort. Parabiosis cohort 1 was generated with one young (5 months) and one old (26 months) male mice. Parabiosis cohort 2 was generated with 4-6 young (5 months) and 4-6 old (21 months) male mice. In cohort 1, 11,771 high quality transcriptomes were collected, using one SVZ sample per 10x lane. In cohort 2, 13,824 high quality transcriptomes were collected, using lipid-modified oligonucleotides to multiplex SVZ samples across three 10x lanes. UMAP projection and cell type clustering of SVZ single cell transcriptomes in cohorts 1 and 2. Each dot represents the transcriptome of an individual cell. Colored by age and intervention (heterochronic parabiosis). For UMAP colored by cell type, see Extended Data Fig. 4a. **b,** Density plots of the predicted chronological ages for aNSC-NPCs from cohort 1 and cohort 2. Green arrows illustrate the median shift in predicted age between old aNSC-NPCs exposed to young blood (heterochronic old) compared to old exposed to old blood (isochronic old control). Density plots for individual mice are provided on the right. **c,** Summary of heterochronic parabiosis effects on chronological age scores across cell types. Effect sizes are calculated by taking the difference in median predicted ages between conditions. Results from both BootstrapCell and EnsembleCell preprocessing and prediction methods are shown for robustness. **d,** Density plots of the predicted biological age scores for neuroblasts from cohort 1 and cohort 2. Green arrows illustrate the median shift in predicted age between old neuroblasts exposed to young circulation (heterochronic old) compared to old exposed to old circulation (isochronic old control). Density plots for individual mice are provided on the right. **e,** Summary of heterochronic parabiosis effects on biological age scores across cell types. Effect sizes are calculated by taking the difference in median predicted ages between conditions. Results from both BootstrapCell and EnsembleCell preprocessing and prediction methods are shown for robustness.

Applying our suite of aging clocks, we predicted both the chronological and biological ages of different cell types in response to heterochronic parabiosis. Interestingly, exposure to young blood had a striking rejuvenation effect on aNSC-NPCs, across both cohorts and for chronological age (rejuvenation of 5.38 months in cohort 1 and 3.66 months in cohort 2; 4.52 months, averaging both cohorts, Fig. 3b, c, Extended Data Fig. 4b-d) and biological age (rejuvenation of 3.57 months in cohort 1 and 1.44 months in cohort 2; 2.51 months, averaging both cohorts) Fig. 3d, e, Extended Data Fig. 4e-g). In other cell types, there was a tendency towards a rejuvenation effect, particularly in neuroblasts and microglia (though with less consistency between cohorts in microglia) (Fig 3c-e, Extended Data Fig. 4b, c, e, f). Overall, the first cohort (21 months difference between young and old) showed a stronger rejuvenation effect than the second cohort (15.5 months difference between young and old) (Fig. 3c, e), suggesting an improved effect of exposure to young blood on older animals (or a greater magnitude when the difference in age in parabionts is larger). Conversely, aging clocks also revealed that young mice exposed to old blood experienced an increase of predicted chronological age across nearly all cell types (Fig. 3c, e), confirming the detrimental impact of old blood in other tissues^70,74–78^.

Thus, single cell-based aging clocks, even when trained on chronological age, can be used to quantify the impact and magnitude of rejuvenation and pro-aging interventions on different cell types. Furthermore, these clocks uncover strong cell-specific rejuvenating effect in old neurogenic regions exposed to young blood focused on proliferating NSCs.

### Exercise rejuvenates cell-specific aging clocks, notably oligodendrocyte clocks

We asked if other interventions that are beneficial for health had a similar or different effect than parabiosis on our suite of cell-type-specific aging clocks. Thus, we applied clocks to another systemic intervention – exercise. Exercise via voluntary wheel running has beneficial effects on the brain, increasing hippocampal neurogenesis and improving memory^79–81^, though the effects of exercise on the SVZ neurogenesis remain under debate^82–84^. We had young and old mice exercise by providing 5 weeks of access to freely spinning wheels (or no wheels as controls) and verified that mice with this paradigm exercised (Fig. 4a, Extended Data Fig. 5a, Supplementary Table 1). We then generated 79,488 single cell transcriptomes from the SVZ neurogenic region from young (5 month) and old (23 month), exercised and non-exercised controls – a total of 15 mice (Fig. 4a) (see Methods). These data represent a great resource for the young and old SVZ neurogenic niche response to exercise.

**Figure 4:**
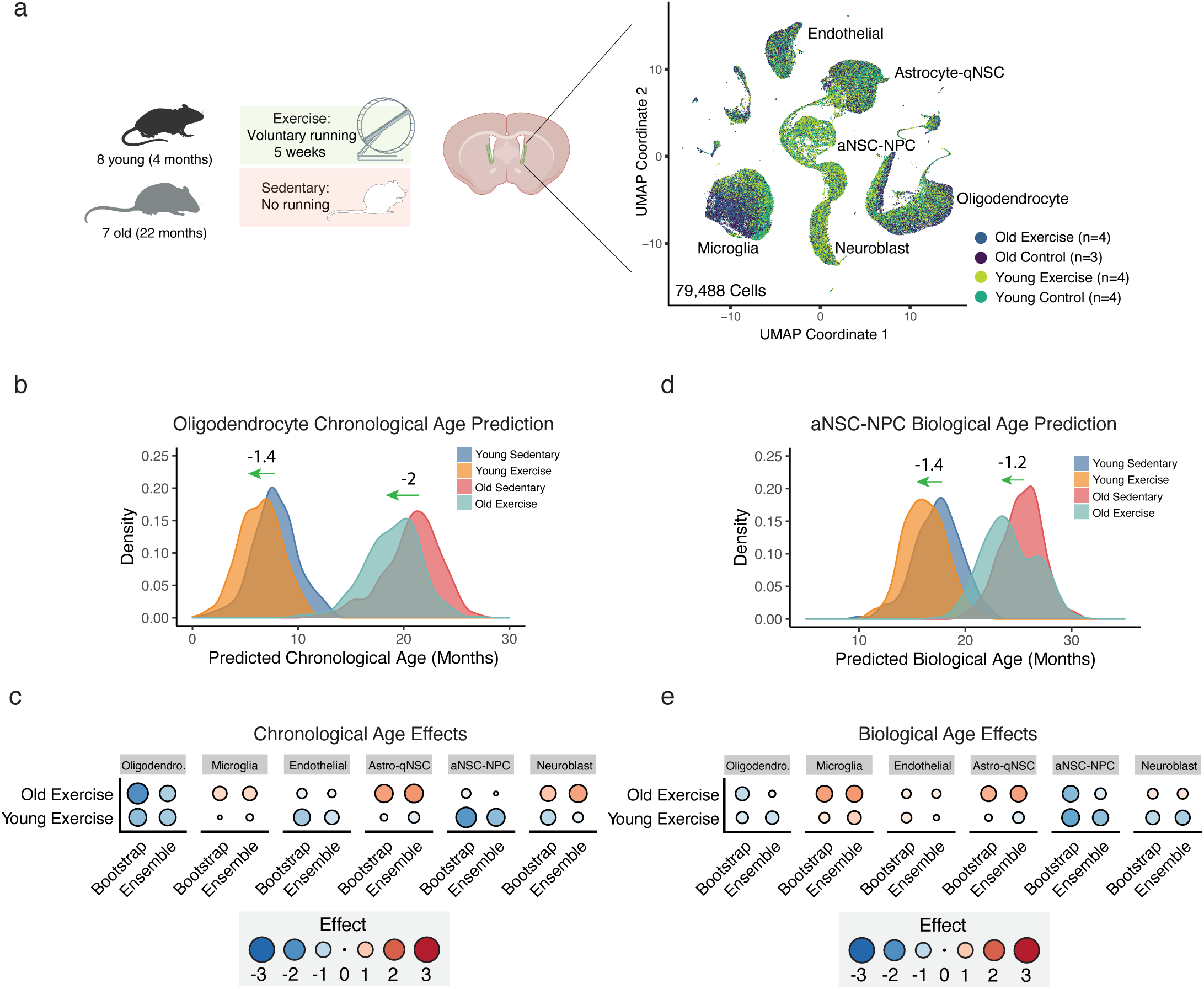
Effect of exercise on cell-type specific aging clocks. **a,** Schematic of voluntary wheel running experiment and UMAP projection of single cell transcriptomes. For the exercise cohort, 4 young (4 months) or 3-4 old (22 months) male mice were transferred into cages with either a freely spinning wheel or no wheel. Wheel rotations were tracked to verify that mice indeed exercised. After 5 weeks, SVZs were collected and 15 lanes of 10x Genomics transcriptomics performed without sample multiplexing. UMAP projection and cell type clustering of SVZ single cell transcriptomes in the exercise cohort. Each dot represents the transcriptome of an individual cell. Colored by age and intervention (exercise). For UMAP colored by cell type, see Extended Data Fig. 5a. **b,** Density plot of predicted chronological ages of oligodendrocytes by age and exercise condition. Exercise consistently rejuvenates oligodendrocyte transcriptomes regardless of age. **c,** Summary of exercise effects on chronological age scores across cell types and ages. Effect sizes are calculated by taking the difference in median predicted ages between conditions. Results from both BootstrapCell and EnsembleCell preprocessing and prediction methods are shown for robustness. **d,** Density plot of aNSC-NPC predicted biological ages. Exercise rejuvenate aNSC-NPC transcriptomes of both young and old mice. **e,** Summary of exercise effects on biological age scores across cell types and ages. Effect sizes are calculated by taking the difference in median predicted ages between conditions. Results from both BootstrapCell and EnsembleCell preprocessing and prediction methods are shown for robustness.

Applying our chronological and biological aging clocks to the exercise transcriptome dataset revealed that exercise led to a consistent decrease of chronological age in oligodendrocytes (1.4 months in young, 2.0 months in old) (Fig. 4b, c) and a small rejuvenation in aNSC-NPCs (1.9 months in young, 0.3 months in old) (Extended Data Fig. 5b). The rejuvenation effect in oligodendrocytes and aNSC-NPCs was also captured using the biological clock (0.6 months in young, 0.8 months in old and 1.4 months in young, 1.2 months in old) (Fig. 4d, e, Extended Data Fig. 5c). Hence, single cell-based aging clocks unbiasedly identify a rejuvenating effect for exercise in old neurogenic regions, notably in oligodendrocytes and aNSC-NPCs, suggesting a selective effect of exercise on some cell types.

### Comparison between exercise and heterochronic parabiosis

We compared the effect of heterochronic parabiosis and exercise. Overall, heterochronic parabiosis had a larger effect than exercise across cell types (Fig. 5a, Extended Data Fig. 6a, c), suggesting that exposure to young blood may be a stronger intervention, at least at the transcriptomic level. Cells that were most responsive to the interventions also differed: aNSC-NPCs were most responsive to heterochronic parabiosis whereas oligodendrocytes (and to a lesser degree aNSC-NPCs) were most responsive to exercise (Fig. 5a, Extended Data Fig. 6a).

**Figure 5:**
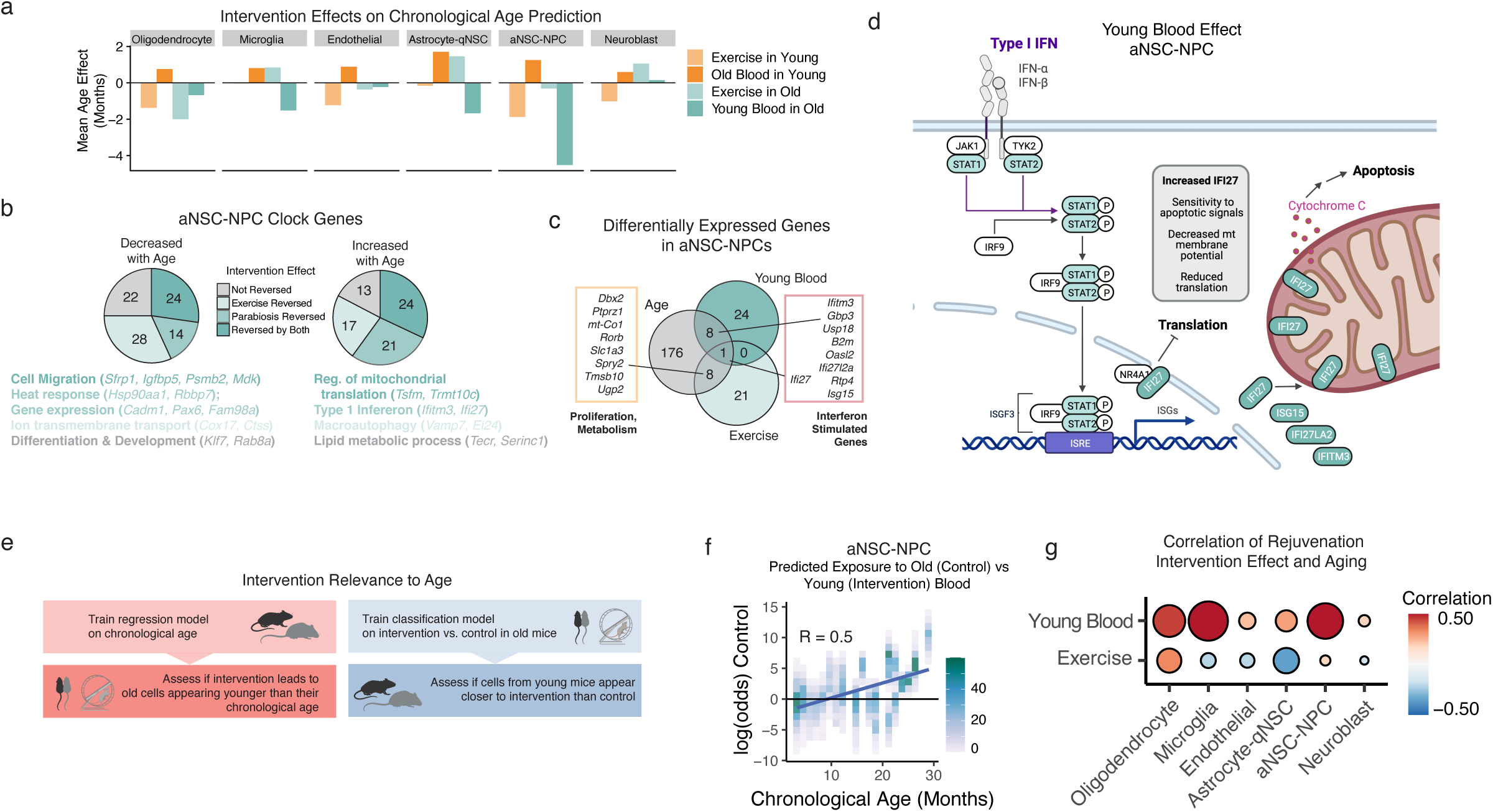
Comparison of exercise and parabiosis interventions on cell-type specific aging clocks reveals distinct modes of action. **a,** Barplot comparing effects of different interventions. Bar represents the difference (in months) between predicted chronological ages between controls and intervention. Parabiosis cohorts 1 and 2 were averaged. **b,** Pie charts of the directional impact and overlap of interventions effects on aNSC-NPC chronological aging clock genes (BootstrapCell). Top Biological Process terms and representative genes are listed underneath. **c,** Overlap of aging and intervention differentially expressed genes in aNSC-NPCs. Differential expression thresholds required a minimum 10% expression change with FDR < 0.1. Differentially expressed genes (DEGs) shared between aging and young blood were interferon- stimulated genes. DEGs shared between aging and exercise were genes involved in proliferation, metabolism, and development. **d,** Schematic of Type I interferon pathway. Exposure to young blood reverses the expression of interferon-stimulated genes. IFN: Interferon; ISGF3: Interferon-stimulated gene factor 3; IRF9: Interferon regulatory factor 9; ISRE: Interferon-sensitive response element. **e,** Predicting ‘rejuvenation intervention or control’ state on the transcriptomes from mice of different ages to assess intervention relevance to aging. **f,** Density scatterplot of logistic regression parabiosis state classification results (log(p(Control)/p(Intervention)) given aNSC-NPC BootstrapCell transcriptomes from mice of different ages. Old mice are more likely to be classified as ‘isochronic old control’ whereas young mice are more likely to be classified as ‘heterochronic old’, indicating that the gene signature that distinguishes exposure to young and old blood is relevant to aging. Higher log(odds) indicates transcriptomes are less likely to be from the rejuvenation intervention (heterochronic old exposed to young blood) and more likely to be from intervention control (isochronic old). R is Pearson correlation. Higher correlation indicates that the main intervention signature overlaps with and reverses age-related changes. **g,** Summary of correlations between intervention state prediction and chronological age across cell types and interventions, with a separate classifier built for each. The exercise classifiers were built to distinguish old sedentary from old exercised transcriptomes for each cell type. The lower correlations between intervention state predictions and age for the exercise samples implies that the signatures that distinguishes exercised and sedentary mice are less related to aging than those derived from parabiosis intervention classifiers. Larger positive correlations are observed in microglia and aNSC-NPCs indicating that the effects of young blood exposure to these cell types are particularly relevant to transcriptomic aging reversal.

We directly compared the genes responding to either or both of these interventions in a cell type responding to both interventions (aNSC-NPCs) (Fig. 5b). Clock genes in aNSC-NPCs that responded to heterochronic parabiosis were mostly genes that increased in expression with aging (such as those associated with a type I interferon response) (Fig. 5b, right panel, Extended Data Fig. 6b, bottom panel). In contrast, clock genes in aNSC-NPCs that responded to exercise were mostly genes that decreased in expression with aging (such as those associated with transmembrane transport) (Fig. 5b, left panel, Extended Data Fig. 6b, top panel). We also examined the overlap between differentially expressed genes in aNSC-NPCs as a function of age and in response to parabiosis and exercise (see Methods). There was minimal overlap between parabiosis- and exercise-responsive genes, suggesting that these interventions impact the aging transcriptome through different mechanisms (Fig. 5c). Young blood specifically reduced interferon stimulated genes (including the shared gene *Ifi27*) (Fig. 5c, d). Exercise actually increased some inflammation genes (*Ifi27*) (Fig. 5c), but reversed the age-associated decline of several genes involved in proliferation and neurogenesis, including *Dbx2* which is implicated in age-related SVZ neurogenic decline^85^ (Fig. 5c).

Finally, we determined whether the main effects of exposure to young blood and exercise are indeed relevant to aging (Fig. 5e). To this end, instead of training regression clocks on age to compare rejuvenation interventions, we trained classifiers on rejuvenation interventions (‘rejuvenated’ and ‘control’) and determined if chronological ages were classified as rejuvenated or control. With the classifier built on heterochronic parabiosis, younger mice showed a greater likelihood of being classified as ‘rejuvenated’ whereas older mice showed a greater likelihood of being classified as ‘control’ (Fig. 5f). This effect was particularly strong in aNSC-NPCs and microglia (Fig. 5f, g). In contrast, with the classifier built on exercise, younger mice showed a greater likelihood of being classified as ‘rejuvenated’ in only two of six cell types (aNSC-NPCs and oligodendrocytes) (Fig. 5g), the same two cell types that showed the strongest rejuvenation from aging clock analysis (see Fig. 5a). This analysis indicates that exercise and young blood induce changes that are indeed relevant to aging and corroborates the comparatively larger effects of young blood as an intervention. Collectively, these machine-learning analyses show cell-type and gene specificity for rejuvenation interventions.

## Discussion

Here we show that single cell RNA-seq data allow the generation of quantitative aging clocks that can be trained on chronological age or on aspect of tissue fitness – i.e. proliferative fraction of stem cells in the neurogenic region. To our knowledge, these are the first quantitative aging clocks based on single cell RNA-seq. We also generate three datasets that represent valuable stand-alone resources: a high temporal resolution single cell RNA-seq aging dataset of a neurogenic niche and the first single cell RNA-seq datasets for a neurogenic niche following heterochronic parabiosis and voluntary exercise. These datasets will be helpful to identify additional cellular and molecular changes during aging and rejuvenation.

Our clock accuracy (e.g. R = 0.92 in microglia) approaches that of bulk DNA methylation and proteomics^13,17,31^ while preserving cell-type-specificity and avoiding biased sorting procedures. Single-cell DNA methylation and proteomic methods have suffered from sparsity and scaling challenges, though there is rapid innovation to address these issues^35,86^. While the methods described here preserve cell-type specificity without relying on cell sorting, ‘pure’ single-cell trained clocks were not as effective as our BootstrapCell and EnsembleCell approaches (see Fig. 1). Thus, using small pools of 15 single cell transcriptomes can mitigate some technological (e.g. gene dropouts) or biological (e.g. transcriptional bursting) challenges inherent to single cell RNA-seq datasets. Nevertheless, increased gene variability (transcriptional noise) is itself a feature of aging^87–91^, and it will be important to model this feature in the next-generation aging clocks.

A limitation of the application of chronological-age-trained clocks is that interventions that stimulate age-associated compensatory pathways (e.g. stress responses) will reflect as age-acceleration despite their functional benefit to the cell, the tissue, or the organism. Thus, there is a need for a better understanding of genes contributing to the aging clocks and their function as well as continued development of functional and phenotypic-trained models. Here, we built clocks based on a functional phenotype of the regenerative region (neural stem cell proliferative capacity), but more comprehensive phenotyping approaches will be important to pursue. Overall, functional-aging clocks are likely to prove instrumental to understanding the biology of aging and rapidly evaluating interventions necessary to extend healthy lifespan.

The observation that heterochronic parabiosis and exercise can ‘turn back’ the single cell-based aging clocks provides a proof-of-concept that these aging clocks, even when trained on chronological age, can record aspects of aging biology. This is in line with other aging clocks built on bulk datasets^13,22–25,32,33^. Our results also highlight cell type specificity for aging and rejuvenation interventions, with some cell types being more responsive than others (e.g. NSCs, oligodendrocytes). This is unique to single cell-based clocks and will allow a better understanding of cell heterogeneity in tissue aging and rejuvenation. Our data also reveal different potential for rejuvenation strategies, at least at the transcriptional level, and different cellular and molecular mechanisms of action. For example, young blood (heterochronic parabiosis) has a stronger rejuvenating effect than exercise, especially in aNSC-NPCs. These results raise the exciting possibility that aging clocks can serve to rapidly test the efficacy of rejuvenation interventions and to support combining specific interventions to counter aging and age-related diseases.

## Acknowledgements

We thank members of the Brunet lab, especially Claire Bedbrook, Paloma Navarro, and Robin Yeo for reading the manuscript and critical discussion. We thank Claire Bedbrook, Paloma Navarro, and Param Priya Singh for independent checking of the code. We thank Christopher S. McGinnis from the Gartner Lab at the University of California, San Francisco for providing MULTI-seq reagents. We thank Anshul Kundaje for fruitful discussion. We thank Liana Bonanno and Jian Luo for conducting surgeries in Parabiosis cohort 2, and Andrew Yang and Tal Iram from the Wyss-Coray lab for facilitating SVZ collection in Parabiosis cohort 2. We thank the Stanford Shared FACS Facility for FACS use and technical support. Elements of figures were created with BioRender.com. Supported by P01AG036695 (A.B., T.A.R, M.A.G.), Chan Zuckerberg Initiative award (A.B.), Simons Foundation grant (A.B.), R01AG072255 (T.W.C), and a generous gift from Michele and Timothy Barakett (A.B.).

## Authors contributions

M.T.B. and A.B. planned the study. M.T.B. generated all datasets and performed all analyses, except for those indicated below. E.S. performed analyses with EnsembleCell clocks, some of the external clock validation and gene overlap, and independent code checking. B.M.G. and N.S. performed heterochronic parabiosis pairings, under the supervision of I.L.W. and T.W.C., respectively. L.L. set up the exercise intervention under the supervision of T.A.R. J.M.R and L.L. provided help in the analysis of the exercise data, under the supervision of M.A.G. and T.A.R., respectively. L.X. helped with dissection of neurogenic regions for single cell RNA-seq and independent code checking. M.T.B. and A.B. wrote the manuscript, and all authors provided comments.

## Data availability

All raw sequencing reads and key processed files is accessible at BioProject PRJNA795276.

## Code availability

The code used to analyze genomic data in the current study are available in the Github repository for this paper (https://github.com/sunericd/svz_singlecell_aging_clocks).

## Methods

### Animals

For aging cohorts and exercise cohort, mice used were male C57BL/6 mice obtained from the NIA Aged Rodent colony. For parabiosis cohort 1, old mice were male C57BL/6 mice from the NIA Aged Rodent colony and young mice were male B6.SJL-*Ptprc^a^ Pepc^b^*/BoyJ male (Pep boy) from the Jackson Lab. For parabiosis cohort 2, old mice were male C57BL/6J and young mice were male C57BL/6J or C57BL/6-Tg(UBC-GFP)30Scha/J from The Jackson Laboratory. At Mice were housed in the Comparative Medicine Pavilion, ChemH/Neuroscience Vivarium, or the SIM-1 Non-Barrier Rodent Facility at Stanford, or in the Veterinary Medical Unit at the Palo Alto VA. All these facilities provide equivalent standard conditions with 12-hour light/dark cycle and ad libitum food and water. All mice were acclimated to their vivarium for at least 2 weeks prior to use in any experiment and care was overseen by the Veterinary Service Center at Stanford University under IACUC protocol 8661.

### Tissue Collection

SVZ neurogenic niches were collected and processed as described in^1^. Briefly, mice were sedated with 1 ml of 2.5% v/v Avertin (Sigma-Aldrich, T48402-25G) and perfused with 15 ml of PBS (Corning, 21040cv) with heparin sodium salt (50 U ml^−1^) (Sigma-Aldrich, H3149-50KU) to remove the blood, and brain collection was performed immediately. As previously described^2^, the SVZ from each hemisphere was micro-dissected and dissociated with enzymatic digestion with papain at a concentration of 14 U ml^−1^, rocking for 10 min at 37°C. The dissociated SVZ was then triturated in a solution containing 0.7 mg ml^−1^ ovomucoid and 0.5 mg ml^−1^ DNaseI (Sigma-Aldrich, DN25-100MG) in DMEM/F12 (Thermo Fisher, 11330032). The dissociated cells from the SVZ were then centrifuged through 22% Percoll (Sigma-Aldrich, GE17-0891-01) in PBS to remove myelin debris. After centrifugation, cells were filtered through a 35-μm snap-cap filter (Corning, 352235), washed once with 1.5 ml of FACS buffer (HBSS (Thermo-Fisher, 14175103), 1% bovine serum albumin (Sigma, A7979), 0.1% glucose (Sigma-Aldrich, G7021-1KG) and spun down for 5 min at 300g. Cells were then resuspended in 120 μl FACS buffer with live/dead staining was performed using 1 μg ml^−1^ propidium iodide (BioLegend, 421301) and kept on ice until sorting. FACS sorting was performed on a BD FACS Aria II sorter, using a 100-μm nozzle at 13.1 PSI. Cells were sorted into low protein binding microcentrifuge tubes containing 750 μl of PBS with 1% BSA and 0.1% glucose. When not applying sample multiplexing (Parabiosis Cohort 1 and Exercise mice), cells were then centrifuged (300g for 5 min at 4°C) and resuspended in 50 μl FACS buffer, counted, and then immediately run on 10x Chromium to capture single cell transcriptomes.

### Lipid Modified Oligonucleotide (LMO) Multiplexing

Sample multiplexing was performed using lipid-modified oligonucleotides (LMOs), a method also known as MULTI-seq^3^. Lipid anchor and co-anchor reagents were kindly provided by the Gartner Lab at the University of California, San Francisco and custom oligonucleotides were ordered from Integrated DNA Technologies. We followed the exact protocol outlined by McGinnis et al (2019)^3^ with the following modifications: (1) All labelling with LMOs was performed in a 4°C cold room because we noticed that the quality of labelling was very sensitive to temperature; (2) To avoid cell loss and cell clumping, cells were sorted into PBS with 2% BSA. BSA was removed through 3 PBS washes; (3) Concentrations and volumes were adjusted to account for low cell numbers: 7.5 μl of 1 mM lipid anchor with oligonucleotide barcode mix was added to a 70 μl volume of resuspended cells followed by 7.5 μl of 1 mM lipid co-anchor; (4) Labelling reactions were quenched with 2% BSA then samples were pooled prior to subsequent 1% BSA PBS washes to further reduce cell loss. The combined sample was ultimately resuspended at 50 μl for cell counting and single-cell RNA-sequencing.

### Single-cell Libraries and RNA-sequencing

Single-cell RNA-sequencing was performed using a 10x Chromium machine and 10x Genomics V3.0 Transcriptomics kits (aging cohorts, parabiosis cohort 2, and exercise cohort) or 10x Genomics V2 kit (parabiosis cohort 1). For sequencing, 10,000 cells per lane were targeted but typical yields were approximately 5,000 cells. Library preparation was done according to the manufacturer’s protocol (10x Genomics V3.0 or 10x Genomics V2 for parabiosis cohort 1). Sequencing was done to target a minimum of 25,000 reads/cell for transcriptome characterization and 5,000 reads per cell for LMO label recovery. The aging cohorts and the parabiosis cohort 2 samples were multiplexed with 4-8 samples per 10x Chromium lane. The parabiosis cohort 1 and the exercise samples were not multiplexed with LMO reagents. Sequencing was performed on either an Illumina HiSeq 4000 (aging cohorts and parabiosis cohort 1) or NovoSeq in 2×150bp setting (parabiosis cohort 2 and exercise).

### Analysis (QC)

CellRanger (version 3.0.2) default settings were used to distinguish cells from background. Subsequent analysis was performed using R (version 3.6.3). Cells were filtered out in Seurat (version 3.2.3)^4^ if they contained fewer than 500 genes or greater than 10% mitochondrial reads. Small clusters of doublets that shared several marker genes from pure populations were identified and removed. LMO demultiplexing was performed using Seurat’s *HTODemux* function. A complete view of the data processing and QC parameters can be found at https://github.com/sunericd/svz_singlecell_aging_clocks.

### Cell Type Annotation

Cell types in all datasets were manually annotated as in^1^ and cross-referenced with annotations present in single cell database PangaloDB^5^. Cell cycle annotation (G1, S, G2/M) was done using Seurat’s *CellCycleScoring* function with default parameters.

### Age Prediction

Chronological or biological age were regressed onto log-normalized gene expression values *ln((gene transcripts / cell transcripts)*10,000)* using the R package glmnet (version 4.0.2)^6^. Lasso regression models were used for BootstrapCells and elastic net models were used with EnsembleCell, described separately below. Parameters were optimized with 5-10 fold cross validation. Chronological age was defined as months since birth. Biological age was defined as *35 – (ProliferativeFraction * 100)* where ProliferativeFraction was the number of cells predicted to be in S or G2/M phase divided by the total number of cells from that sample. The number 35 was selected in order to transform biological age into same range as chronological age.

### BootstrapCell

In addition to predicting age from single cell transcriptomes and pseudobulked transcriptomes, we explored alternative ways of preprocessing transcriptomes. To generate a BootstrapCell, 15 single cell transcriptomes were sampled without replacement from the pool of cells of a given cell type from a given animal (e.g. oligodendrocytes from a single mouse). Gene counts were then summed. A number of 15 cells was empirically found to balance the tradeoff between sample number and gene coverage per sample. This bootstrapping process was repeated 100 times for each cell-type-animal combination. BootstrapCells were used as input into lasso regression models. This approach had the effect of normalizing the contribution of each animal rather than each single cell transcriptome.

### EnsembleCell

We devised and evaluated a second preprocessing and age prediction technique to compare to our BootstrapCell approach and to demonstration robustness to changes in preprocessing and model architecture. In the EnsembleCell approach, 20 elastic net models were trained for each cell type. For each model, gene expression data from cells were randomly partitioned into groups of 15 single cell transcriptomes and the unique transcript counts for all cells in each group were summed to create “EnsembleCells”. The unique molecular identifiers (UMIs) were then normalized across samples and log-transformed. An elastic net model was trained to predict the age of the host animal of the cells from the cells’ gene expression profiles. To predict age on new cell gene expression profiles, we used the weighted average of predictions across all 20 models, where weights were determined by the r^2^ of the model on a held-out validation set.

### Heterochronic Parabiosis

Two independent cohorts of heterochronic parabiosis were generated. These parabiosis cohorts were generated in different facilities, by different surgeons, in different years, and they were analyzed with different versions of 10x Genomics single cell transcriptomics kits. Parabiosis cohort 1 involved six mice divided into three pairs. Old parabionts were C57BL/6 male mice from the National Institute on Aging Aged Rodent colony at Charles River. Young parabionts were B6.SJL-*Ptprc^a^ Pepc^b^*/BoyJ male (Pep boy) mice from The Jackson Laboratory and C57BL/6 male mice from the NIA. Of the young, only the Pep boy mice were used for transcriptomics. Congenic (rather than isogenic) pairings were performed to enable verification of blood chimerism by FACS with antibodies specific to CD45.1 (BioLegend 110705) or CD45.2 (BioLegend 109814) alleles. Mice were 4 and 25 month-old at the start of the experiment, and parabiosis was conducted for five weeks until tissue collection, so at collection time mice were 5 and 26 month-old. Pairs were established previously described^7–9^ by suturing the peritoneums of adjacent flanks and joining skin with surgical clips. Five weeks after the parabiosis surgery, mice were anaesthetized with 2.5% v/v Avertin, euthanized by cardiac puncture, and perfused with 15 ml PBS with heparin (50 U ml^−1^). SVZ dissection, digestion, and FACS were performed as describe above. 10x Genomics single cell transcriptome V2 libraries (one sample per 10x lane) were generated and sequenced on one Illumina HiSeq lane by the Stanford Function Genomics Facility. Animal care and parabiosis procedures were performed in accordance with Stanford University under IACUC protocols 8661 and 16246.

Parabiosis Cohort 2 involved 18 male mice divided into four isochronic young, four heterochronic, and six isochronic old. All mice in this cohort were sourced from the Jackson Laboratory and housed in the Veterinary Medical Unit at the Palo Alto VA. Old mice were C57BL/6J and young were C57BL/6J or C57BL/6-Tg(UBC-GFP)30Scha/J. Mice were aged 4 and 19.5 months at the start of the experiment, and parabiosis proceeded five weeks until tissue collection, so at collection time mice were 5 and 21 months old. Surgeries were performed as described above. Five weeks after surgery, mice were anaesthetized with 2.5% v/v Avertin, euthanized by cardiac puncture, and perfused with 15 ml PBS with heparin (50 U ml^−1^). SVZ dissection, digestion, and FACS were performed as describe above. Tissue collection took place on three separate days and samples were multiplexed with lipid-modified oligonucleotides. 10x Genomics single cell transcriptome V3 libraries were generated in house and sequenced by Novogene on an Illumina NovoSeq lane. Animal care and parabiosis procedures were approved VA Palo Alto Committee on Animal Research and listed on ACORP LUO1736.

### Exercise

C57BL/6 male mice sourced from the National Institute on Aging Aged Rodent colony at Charles River at housed in the Veterinary Medical Unit at the Palo Alto VA. Young and old mice were aged 4.5 and 21.5 months at the start of the five week voluntary wheel running intervention. During the intervention period, mice were singly housed in cages accommodating a running wheel. Control mice had no access to a wheel. Running was verified by recording wheel revolutions. After five weeks, mice were anaesthetized with 2.5% v/v Avertin, euthanized by cardiac puncture, perfused, and cell suspensions from dissected SVZs generated as described in above under the *Tissue Collection* section. 10x Genomics V3.0 transcriptomics kits were used to generated libraries without upstream sample multiplexing. Tissue processing occurred across two separate mornings. SVZ libraries were pooled and sequenced on an Illumina NovoSeq.

### Gene Set Enrichment Analysis

Gene set enrichment analysis was executed using Enrichr^10^ to query cell type-specific clock genes for enrichment against GO Biological Process gene sets. Statistics were exported from the Enrichr web tool and processed and visualized in R with ggplot2 (version 3.3.3) package.

### Differential Expression

MAST^11^ software was used to calculate differential expression statistics. To determine age differentially expressed genes, a contrast was made between young mice <7 months old and old mice >20 months old. Permissive 1.1-fold change and 10% FDR cutoffs were applied.

### Intervention Classification Models

To evaluate the aging-relevance of ‘rejuvenation’ interventions, cell type-specific intervention classification models were constructed using cross-validated penalized logistic regression models (*cv.glmnet(type.measure = “mse”, family = “binomial”*) trained on BootstrapCell preprocessed intervention data then applied to aging time course dataset. Positive intervention samples (old exercise, heterochronic old) labeled “0” and control intervention samples (old sedentary, isochronic old) labeled “1”.

---

**Extended Data Figure 1.**
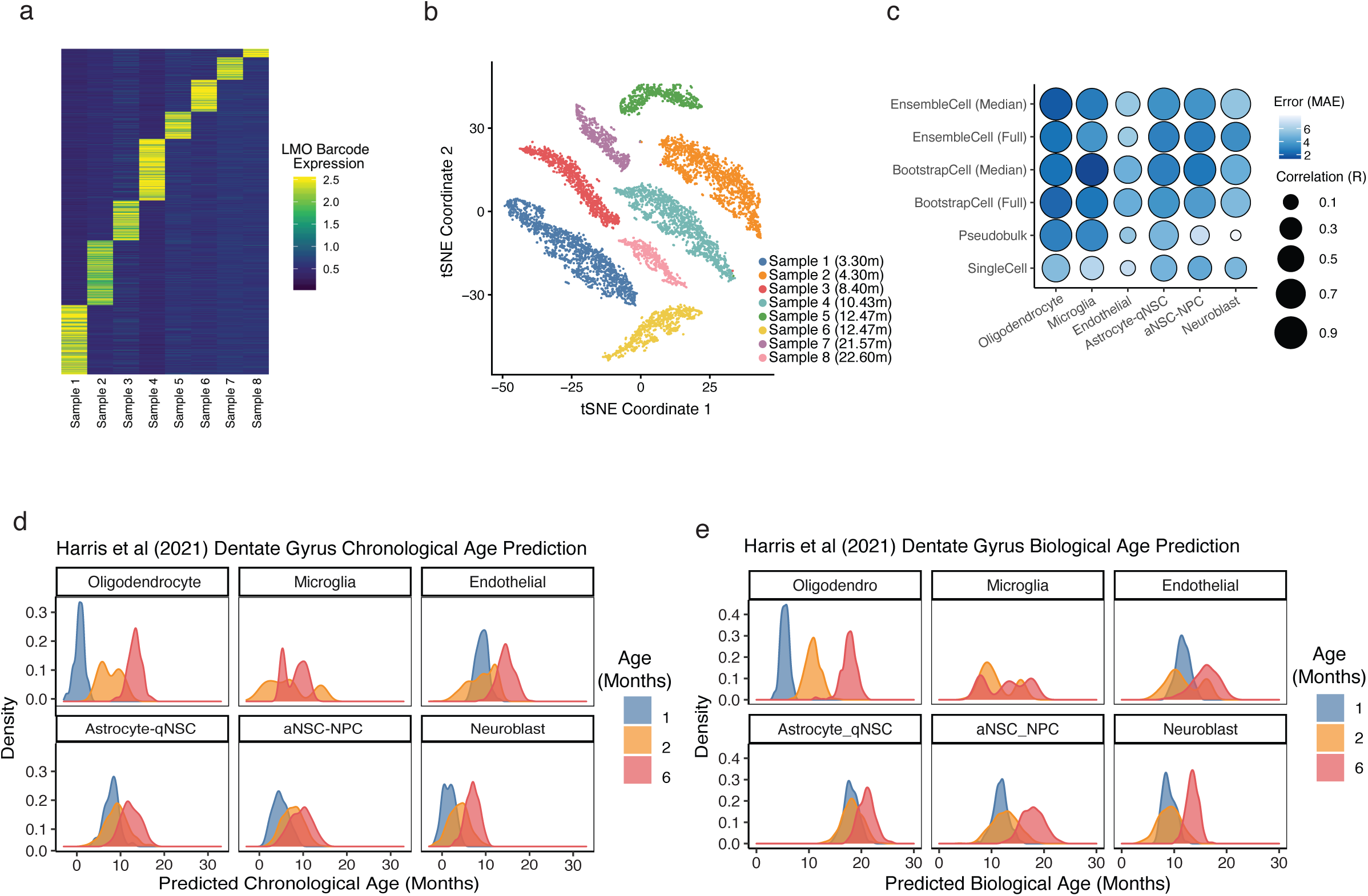
**a,** Lipid-modified oligonucleotide cell barcodes detected from 8 SVZ samples multiplexed in one 10x lane. **b,** Same data as (a) but visualized using tSNE. Note that samples 5 and 6 were from mice of the same age (and were colored in the same color in Fig 1c). **c,** Overview of Pearson correlation coefficients and median absolute error values for tested methods of predicting chronological age across cell types from single-cell transcriptomic data, including both full distribution and median metrics only. Performance is based on cross-cohort-validation. **d,** Application of chronological aging clocks (BootstrapCell) to an external dataset from Harris et al (2021). Transcriptomes of analogous cell types were collected from the hippocampus instead of the subventricular zone. There were no microglia cells in the dataset at the 1 month time point. **e,** Application of biological aging clocks (BootstrapCell) to an external dataset from Harris et al (2021). There were no microglia cells in the dataset at the 1 month time point.

**Extended Data Figure 2.**
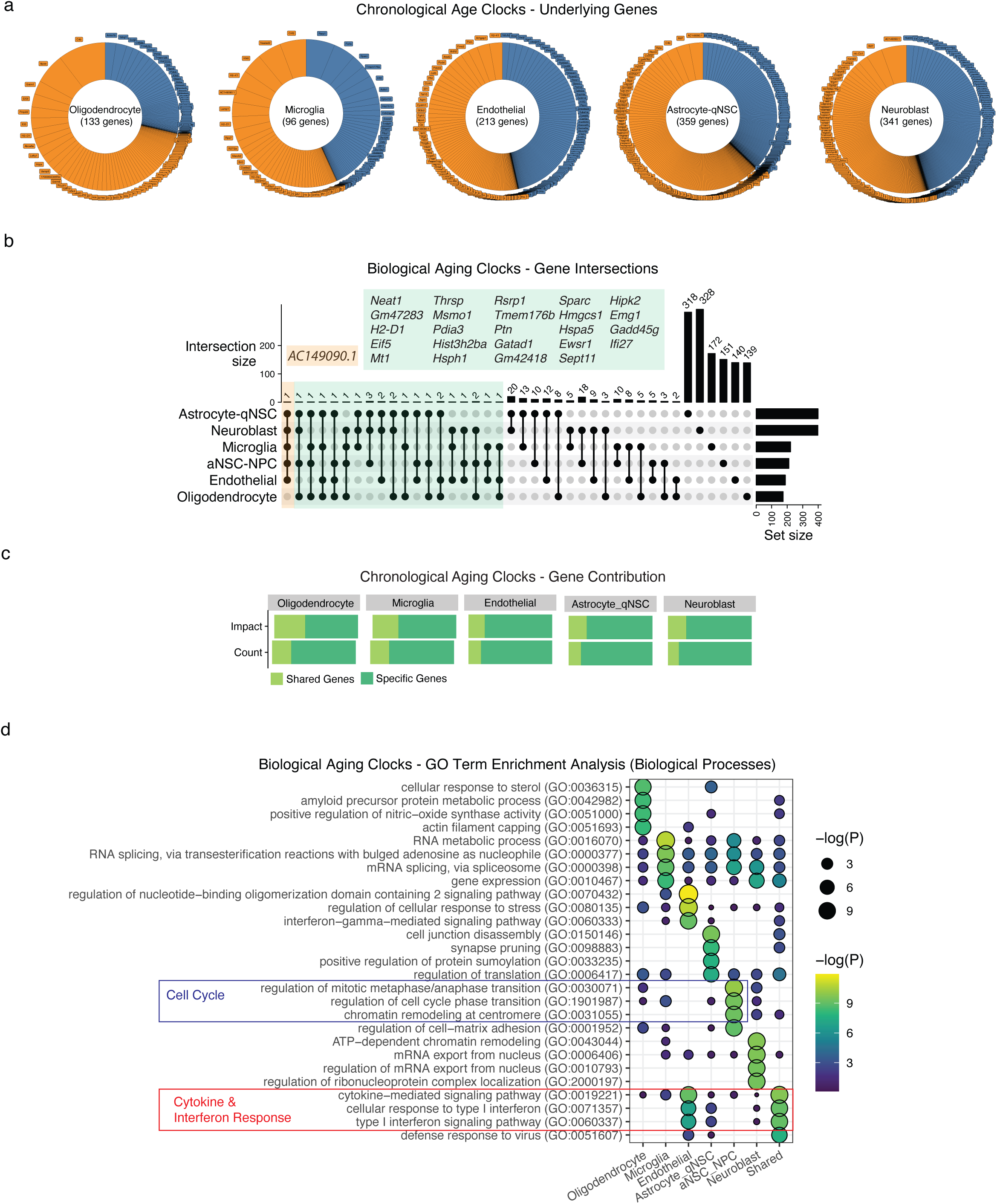
**a,** Genes contributing to chronological aging clocks (BootstrapCell) and their coefficients. Donut plots, with sector size denoting gene weight in the model and color indicating sign of expression change with age. Total number of genes used by the clock is provided in the center of each donut plot. Positive coefficients (orange) indicate higher gene expression is associated with older age. Negative coefficients (blue) indicate lower gene expression is associated with older age. **b,** Upset plot illustrating the intersection of gene sets used by cell type-specific biological aging clocks. No genes were used in all 6 biological aging clocks. **c,** Count and coefficient impact of shared and cell type-specific clock genes for chonological aging clocks. Shared is defined as present in at least one of the other five clocks. **d,** Top enriched Gene Ontology Biological Process terms from gene set enrichment analysis of genes used in biological aging clocks. Shared genes (present in two or more clocks) are enriched for *cytokine-mediated signaling pathway* and *cellular response to type I interferon.* The aNSC-NPC biological aging clock genes are enriched for cell cycle pathways.

**Extended Data Figure 3.**
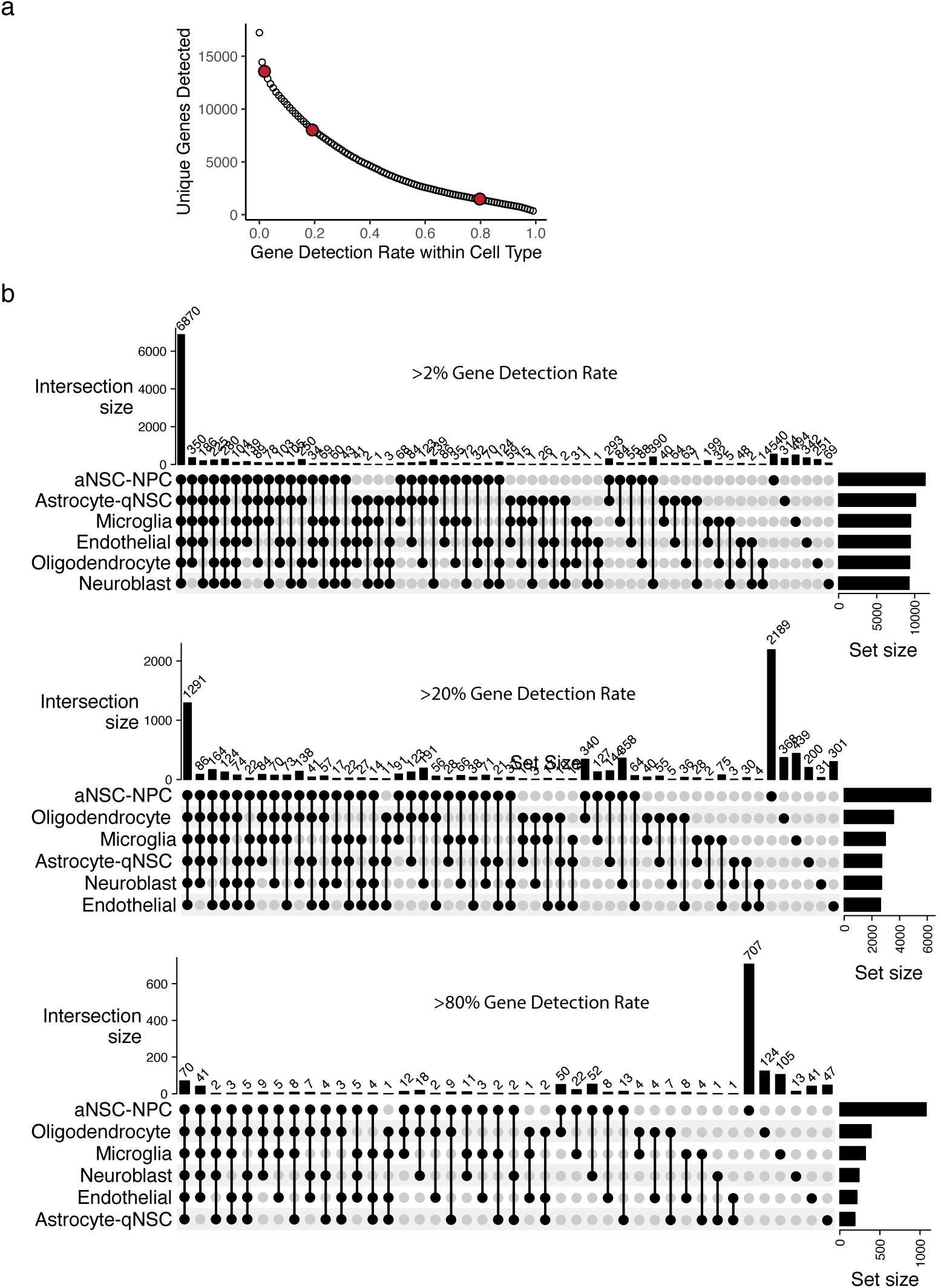
**a,** Unique genes detected in single cell transcriptomes from the subventricular zone as a function of gene detection rate. Red dots indicate unique genes detected at 2%, 20%, and 80% detection rates. **b,** Upset plots showing transcriptome overlaps between cell types at different levels of expression detection. Most genes are shared if a very low threshold of detection is used. Above 80% detection rate, transcriptomes are very cell type specific. However, the shared core of easily detected genes in transcriptomes (70 genes) is much larger than the shared core of genes selected by clocks (∼1).

**Extended Data Figure 4.**
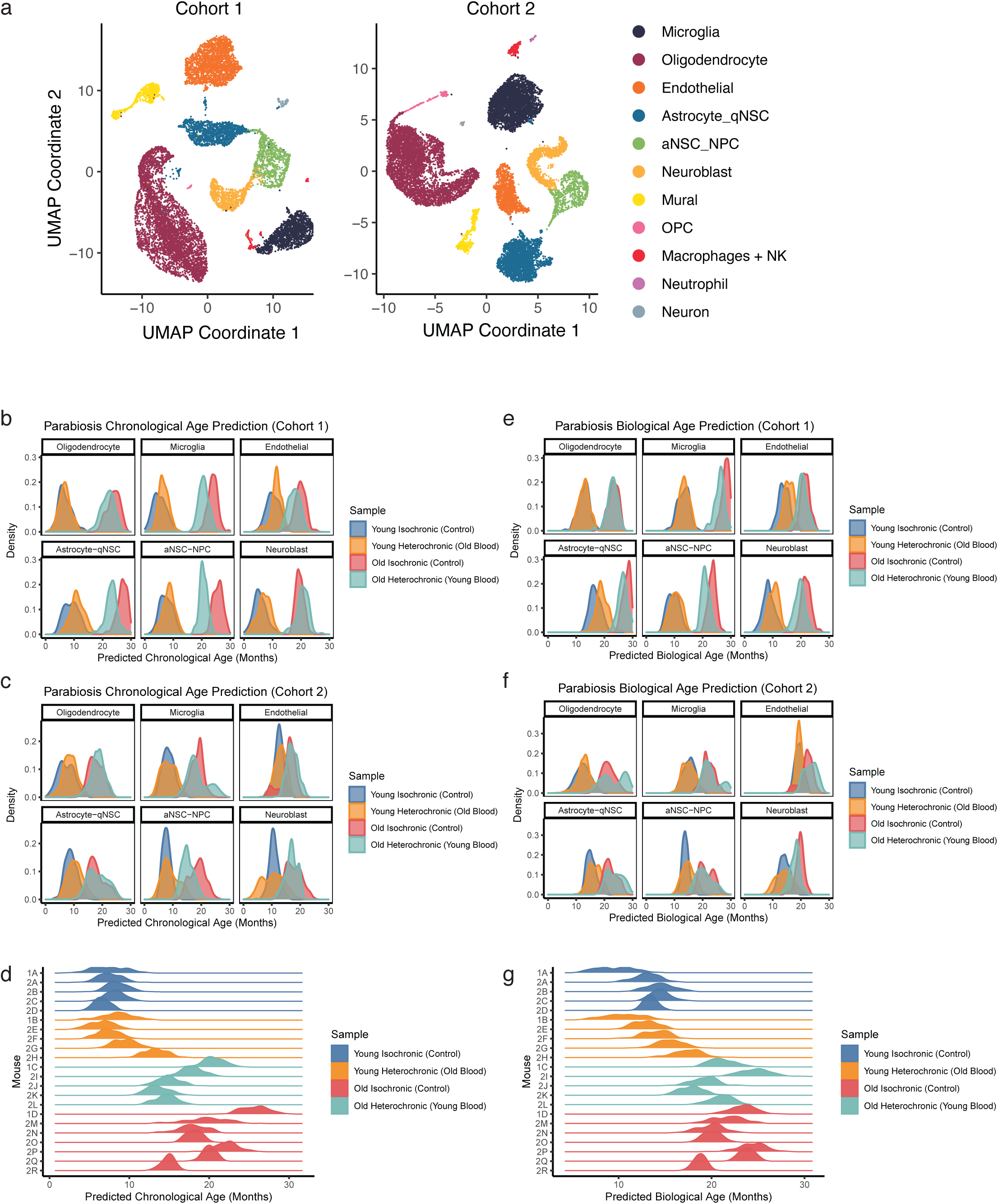
**a,** UMAP projection of single-cell transcriptomes labeled by cell type from both parabiosis cohorts. **b,** Prediction distributions using chronological aging clocks for Parabiosis cohort 1 (BootstrapCell). **c,** Prediction distributions using chronological aging clocks for Parabiosis cohort 2. **d,** Prediction distributions using chronological aging clocks for both cohorts separated by mouse. **e,** Prediction distributions using biological aging clocks for Parabiosis cohort 1. **f,** Prediction distributions using biological aging clocks for Parabiosis cohort 2. **g,** Prediction distributions using biological aging clocks for both cohorts separated by mouse.

**Extended Data Figure 5.**
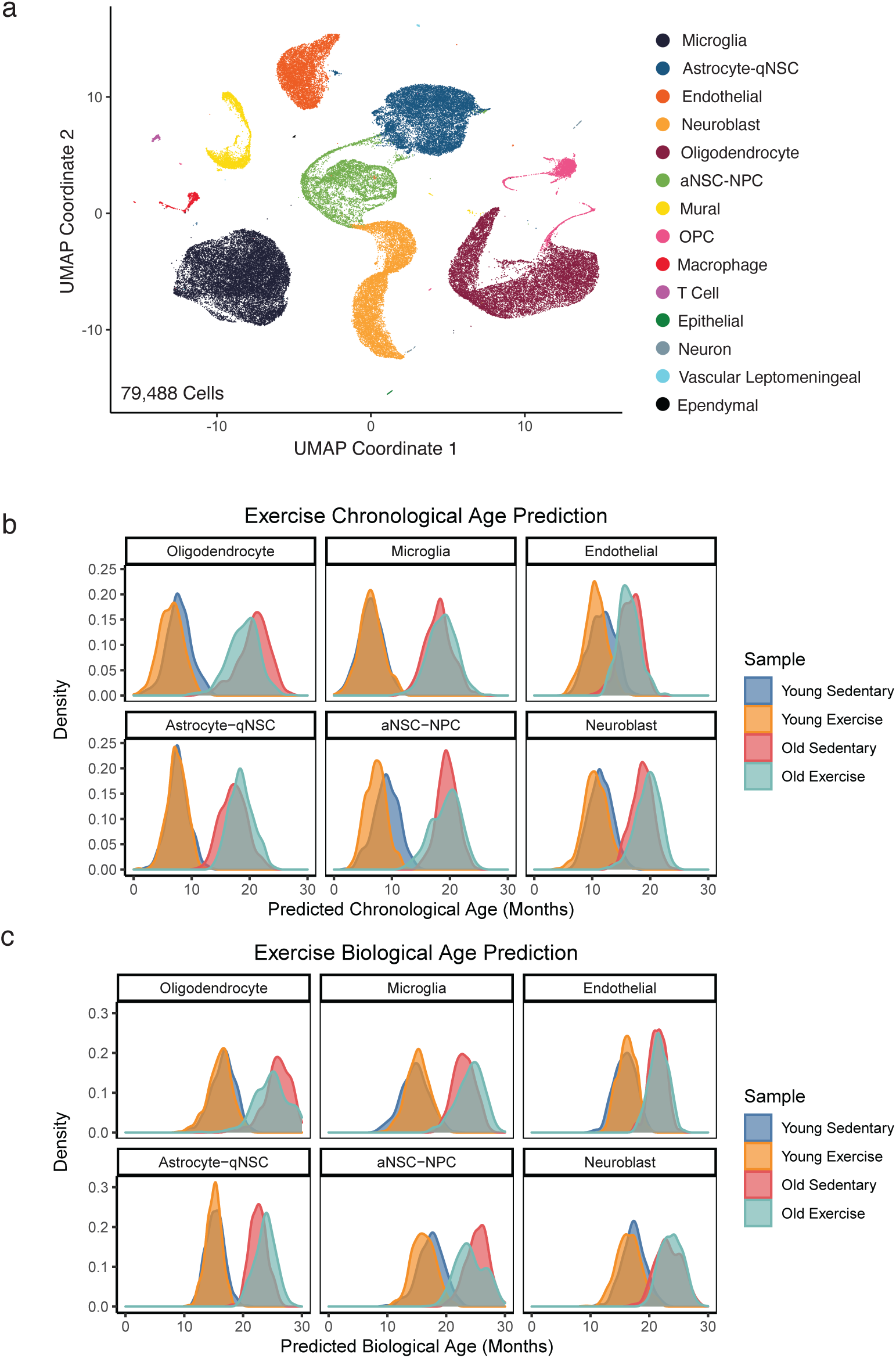
**a,** UMAP projection of single-cell transcriptomes labeled by cell type from the exercise intervention cohort. **b,** Prediction distributions using chronological aging clocks (BootstrapCell) for young and old, exercised and control samples. **c,** Prediction distributions using biological age clocks (Prediction distributions using) for young and old, exercised and control samples.

**Extended Data Figure 6.**
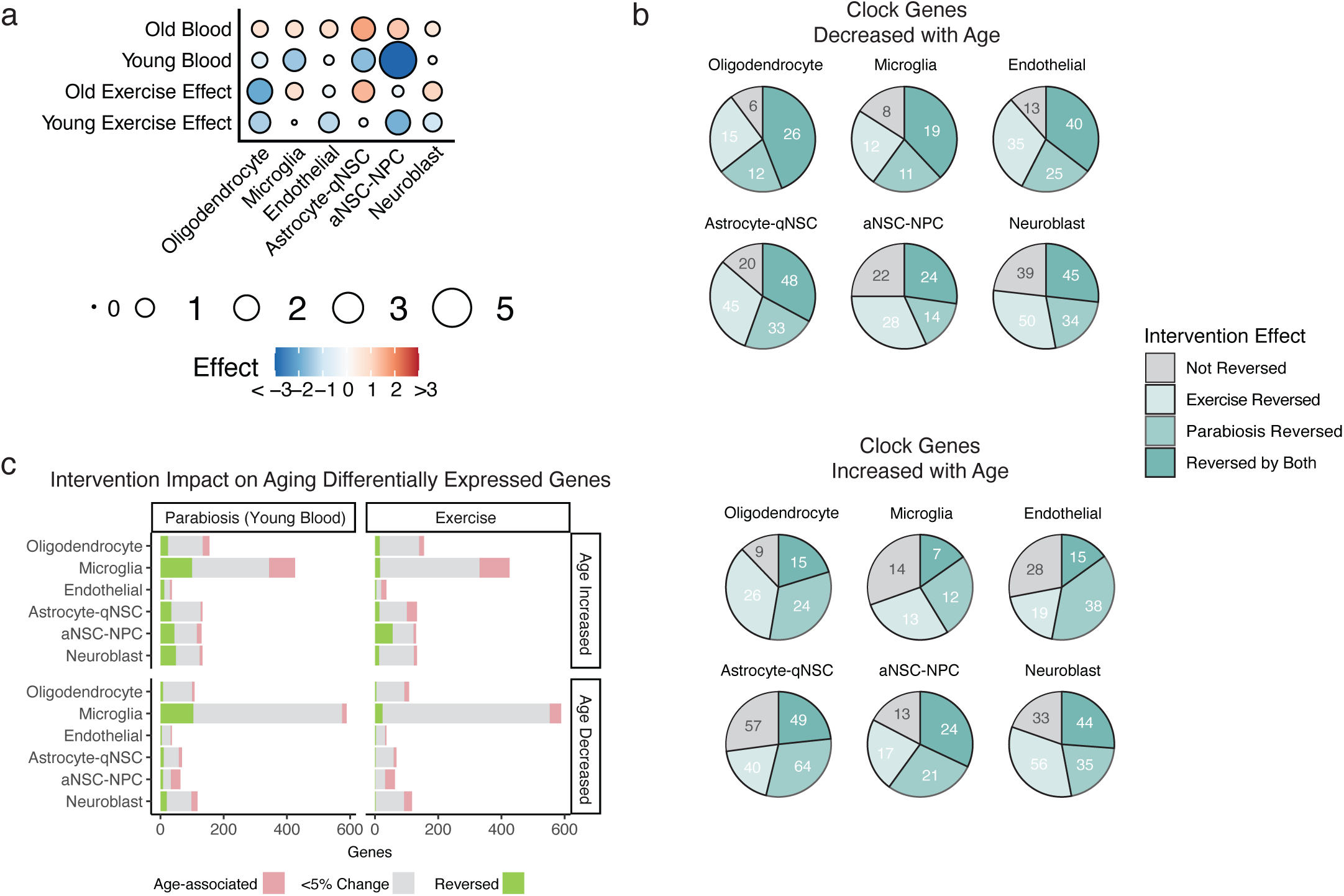
**a,** Dot plot summarizing and comparing intervention effects across cell types. Effect sizes for parabiosis were determined by averaging cohort 1 and cohort 2. Exposure to young blood via heterochronic parabiosis has a stronger rejuvenation effect than exercise, and the impact is strongest in aNSC-NPCs. **b,** Pie charts indicating the overlap and directional effects of different interventions on genes selected by chronological aging clocks (BootstrapCell). Top: selected clock genes increase with age: Bottom: selected clock genes decrease with age. **c,** Barplots showing the proportion of genes that are differentially expressed age which are reversed by intervention, cell type, and whether the genes increase or decrease with age (FDR < 0.1, abs(ln(fold change)) > 0.1, or approximately greater than a 10% change with age). Microglia exhibit the most age-associated changes. Parabiosis is effective at shifting differentially expressed genes during aging towards a more young-associated expression levels (more green in “Parabiosis” column). Reduction of expression of genes that increase with age is larger than the induction of expression of genes that decrease with age (more green in “Age Increased” rows).

## Supplementary Table Legends

**Supplementary Table 1.** Mice metadata including age and multiplexing oligonucleotide sequence.

**Supplementary Table 2.** Prediction performance summary table for all models.

**Supplementary Table 3**. Clock genes and coefficients for all models.

